# Pharyngeal pumping and tissue-specific transgenic P-glycoprotein expression influence macrocyclic lactone susceptibility in *Caenorhabditis elegans*

**DOI:** 10.1101/2020.11.25.397646

**Authors:** Alexander P. Gerhard, Jürgen Krücken, Cedric Neveu, Claude L. Charvet, Abdallah Harmache, Georg von Samson-Himmelstjerna

## Abstract

Macrocyclic lactone (ML) resistance has emerged in many parasitic nematodes including the pathogenic horse roundworm *Parascaris univalens*. The underlying mechanism of ML resistance and the drug penetration routes into the nematodes remain to be elucidated. Drug efflux by P-glycoproteins is considered a potential resistance mechanism but conclusive functional evidence is lacking. To this end, we used a motility assay modified to stimulate pharyngeal pumping (PP) by bacteria or serotonin and tissue-specific expression of *Pun-* PGP-9 in the free-living model nematode *Caenorhabditis elegans*. Here, stimulation of PP was identified as an important factor for *C. elegans* ML susceptibility, increasing the EC_50_ values of ivermectin by up to 11.1-fold and of moxidectin by 1.2-fold. In this context, intestinal *Pun-*PGP-9 expression elicited a protective effect against ivermectin and moxidectin only in the presence of PP stimulation, increasing the EC_50_ values by approximately 3- to 4-fold (ivermectin) or by < 1.3-fold (moxidectin). Conversely, epidermal *Pun-*PGP-9 expression protected against moxidectin regardless of PP with EC_50_ fold changes below 1.5 but against ivermectin with a considerable 2.9-fold EC_50_ increase only when the drug is not actively ingested. Our results highlight the role of active drug ingestion by nematodes for susceptibility and provide conclusive functional evidence for a contribution of P-glycoproteins to ML resistance.

**Author Summary:** Parasitic nematode infections pose a serious threat to animal health, in particular in light of the widespread anthelmintic resistance in different nematode species. In equines, the roundworm *Parascaris univalens* is a major pathogen of foals, exhibiting widespread resistance against macrocyclic lactones (MLs). This represents a particular challenge to animal health, but the underlying mechanisms and drug penetration routes remain mostly unknown. P-glycoprotein ABC-transporters have been linked to ML resistance in several parasitic nematodes. Here we demonstrate by tissue-specific overexpression of *Pun-*PGP-9 in the free-living model nematode *Caenorhabditis elegans* their ability to reduce the susceptibility to two commonly used MLs, ivermectin and moxidectin. At the same time, active drug ingestion by pharyngeal pumping (PP) strongly enhanced ivermectin and moderately effects moxidectin susceptibility. In more detail, the effect of intestinal or epidermal *Pun-*PGP-9 was dependent on active drug ingestion. These observations indicate differences in the drug penetration routes between ML derivatives and allow novel insight into the functional role of P-glycoproteins.

## 1 Introduction

Treatment of parasitic nematode infections in animals and humans relies on chemotherapy. Macrocyclic lactones (MLs) represent the most widely used drug class in veterinary medicine and are considered the best option for many parasitic nematode diseases due to their high efficacy, low toxicity, and broad-spectrum (1). Drug resistance to MLs is widespread in parasitic nematodes of ruminants (2), horses (3), companion animals, (4, 5) and humans (6). With over 1.5 billion infected humans (7) and principally essentially all domestic animals exposed to nematode infections, the ongoing development and spread of ML drug resistance is an obstacle to maintaining health standards. In equines, resistance against MLs and even against multiple anthelmintic classes has emerged in the pathogenic *Parascaris univalens*. The now widespread resistance poses a severe threat to young animals, and resistance is a global challenge (8).

Much of our knowledge regarding the mode of action of MLs in nematodes has been derived from studies using the free-living model nematode *Caenorhabditis elegans*. In *C. elegans* and in parasitic nematodes, the primary pharmacological targets of MLs are the glutamate-gated chloride channels (GluCls), while the ionotropic γ-aminobutyric acid (GABA_A_) receptors are secondary targets (9, 10). Strains carrying multiple mutations in GluCl subunits were shown to be highly resistant to ivermectin (10); however, ML susceptibility can be restored by the transgenic expression of *C. elegans*, and parasite-derived GluCl subunits (11). The irreversible binding of MLs to GluCls leads to a hyperpolarisation of the respective neurons that control locomotion, pharyngeal pumping, and egg laying, resulting in flaccid paralysis of the muscles and pharynx (9, 10). However, how MLs penetrate the worm and reach their target tissue, and the role of metabolism in the resistance mechanism remains mostly unknown. In general, the understanding of the molecular mechanisms of ML resistance in parasitic nematodes has made only moderate progress (12) but recent genome-wide studies have considerably expanded our knowledge of ML resistance mechanisms (13). These studies either point towards a single ivermectin (IVM) resistance locus (14, 15), or a complex multigenic resistance trait (16, 17), depending on the species and isolate.

Since the emergence of ML resistance in the late 1980s, the ATP-binding-cassette subfamily B member 1 (ABCB1) transporter genes, more commonly referred to as P-glycoproteins (Pgp), have been proposed as mediators of anthelmintic resistance in parasitic nematodes (18, 19) and in *C. elegans* (20, 21). These xenobiotic efflux pumps are conserved in eukaryotes and have a broad lipophilic substrate range (22). On the one hand, a role of Pgps in ML resistance in nematodes has been inferred from the observed differences of Pgp expression in association with drug exposure. For example, individual Pgps were found to exhibit constitutively higher expression levels in resistant compared to susceptible worm isolates and genetic signatures of selection were detected in close proximity to Pgp loci in resistant and introgressed populations (16–18, 23). On the other hand, functional studies in *C. elegans* have provided strong evidence of Pgp involvement in the ML resistance phenotype. For example, individual Pgp deletion strains, and silencing or inhibition of Pgps results in increased susceptibility (20, 21). While these studies have shed some light on the putative contribution of Pgp to ML resistance, the functional mechanism how Pgps contribute to ML resistance remains superficial and fragmented.

Many marketed ML derivatives have been shown to exhibit differing pharmacokinetics, efficacies, and chemical properties (1). MLs are classified into two groups, the avermectines (including IVM) and the milbemycines (including moxidectin, MOX), the latter lacking a polysaccharide side chain at C13 of the lactone ring. This chemical disparity has been suggested to lead to the differences found in the affinity of nematode Pgps to different ML derivatives (24, 25). Pharmacokinetics of different ML derivatives are well-studied in mammals, and it was found that they accumulate in fatty tissues due to their high lipophilicity (26). The major route of ML elimination is the intestine and in this process, Pgps have been shown to play an important role (27). In contrast, few studies have explored the uptake routes of MLs in worms and the functional role of Pgp in modulating ML susceptibility. In mammals, Pgps are considered to constitute a barrier to MLs, as animals or humans deficient in the Pgp gene (i.e., *ABCB1* or *mdr-1*) suffer from acute neurotoxicity resulting from MLs crossing the blood-brain barrier (28). In this regard, the capacity of MDR-1 to prevent barrier crossing of MOX and IVM varies between the two derivatives (29).

In contrast to mammals which only possess one (human) to three (rodents) Pgps, the nematode Pgp gene family comprises a diverse repertoire that differs between nematodes species (30). In *C. elegans*, fifteen different Pgp-genes, including a pseudo-gene, have been described (31). More recently, the Pgp repertoire of *P. univalens* was completely deciphered, revealing ten different Pgp genes (32). In the latter study, constitutive expression levels of *Pun-pgp-11.1, Pun-pgp-16.2*, and *Pun-pgp-9* were significantly higher than the other Pgps. Generally, the *pgp-9* gene lineage has been repeatedly associated with ML resistance in several nematode species such as cyathostomins (23, 25), *Teladorsagia circumcincta* (16, 33), and *H. contortus* (34). In general, Pgp expression is most prominent in the intestine in different nematode species, including *P. univalens* (32, 35–37). In addition, the epidermis (= hypodermis) also exhibits moderate Pgp expression levels in *P. univalens* (35) and *C. elegans* (32). Pgp expression in other tissues such as the neuronal or the reproductive system appears at a comparatively low level (32, 35). However, the *in vivo* functional roles such as barrier function, excretion, or other physiological processes of Pgps expressed in specific worm tissues and their potential to reduce ML uptake have never been addressed.

The present study’s objective was to elucidate the details of IVM and MOX uptake and the protective role of *Pun*-PGP-9 at specific uptake barriers. *C. elegans* feed on bacteria which they take up from the environment by pharyngeal pumping (PP) (38). In the absence of an appropriate PP stimulus such as food or 5-hydroxytryptamine (5-HT = serotonin) (39), PP and active ingestion stops (40, 41). In this study, stimulation of PP was used to increase selectively the intestinal exposure to MLs in *C. elegans* by active drug ingestion or to mostly limit exposure to the cuticle-epidermis in the absence of a specific PP stimulus. Concurrently, tissue-specific expression of *Pun*-PGP-9 in the intestine or the epidermis was used to examine the role of Pgp transporter expression at potential ML uptake barriers. Behavioural and pharmacological assays show that IVM and MOX can penetrate *C. elegans* through the intestine and epidermis and that Pgps play a key protective role in reducing worms’ susceptibility to MLs.

## 2 Results

### 2.1 Tissue-specific expression patterns of *Pun*-PGP-9 in a *Cel-pgp*-9 mutant strain

To address the function of *Pun-pgp-9* using *C elegans*, we made use of the available *Cel-pgp-9 (tm830)* null allele. We generated transgenic strains carrying extra-chromosomal arrays giving tissue-specific overexpression of *Pun-pgp-9* in the *Cel-pgp-9* null mutant *(tm830)* background. Two lines henceforth referred to as intestine-Pgp-9 line 1 and line 2 with intestinal *Pun*-PGP-9 expression driven by the intestine specific *gut-esterase-1* (*ges-1)* promotor (42), and pharyngeal GFP expression (*IntPgp-9Ex1 and IntPgp-9Ex2* [*Cel-pgp-9(-); Cel-ges-1p::Pun-pgp-9::FLAG::Cel-unc-54_3′-UTR; Cel-myo-2p::gfp::Cel-unc-54_3’UTR]*), one line henceforth addressed as epidermis-Pgp-9 with *Pun-*PGP-9 expression controlled by the epidermis specific *collagen-19* (*col-19*) promotor (43), and pharyngeal GFP expression (*EpiPgp-9Ex1* [*Cel-pgp-9(-); Cel-col-19p::Pun-pgp-9::FLAG::Cel-unc-54_3′-UTR; Cel-myo-2p::gfp:: Cel-unc-54_3’UTR]*); another line henceforth addressed as the control strain expressing only GFP in the pharynx (*CtrlEx1* [*Cel-pgp-9(-); Cel-myo-2p::gfp:: Cel-unc-54_3′-UTR*).

Transcription of *Pun-pgp-9* was confirmed by RT-PCR for all *Pun*-PGP-9 transgenic strains by amplifying a 1043 bp PCR product while no PCR product was amplified in the control strain (**Fig. 1A**). Protein localisation in the epidermis and the intestine was confirmed by immunofluorescence targeting a FLAG-tag fused to the 3’-end of *Pun*-PGP-9 and visualized using a secondary DyLight405 antibody and confocal microscopy (**Fig. 1B-E**). *Pun*-PGP-9 protein expression in the intestine-Pgp-9 line 1 appeared intestinal specific with expression in all intestinal cells (**Fig. 1B**). At a higher magnification, a strong apical expression is apparent (**Fig. 1E**). Expression in the epidermis-Pgp-9 strain was detectable throughout the hyp7 syncytium and in the anterior epidermis (**Fig. 1C**) while the apical epidermal membrane was stained almost continuously. If fluorescence was visible in both the green and blue channels, e.g., in the representative image of the epidermis-Pgp-9 strain (**Fig. 1C**), it was considered autofluorescence. Specific blue fluorescence was not visible in the control strain following staining, while pharyngeal GFP expression was visible, indicating that the antibody staining was specific to the FLAG-tagged *Pun-*PGP-9 (**Fig. 1D**).

**Figure 1.**
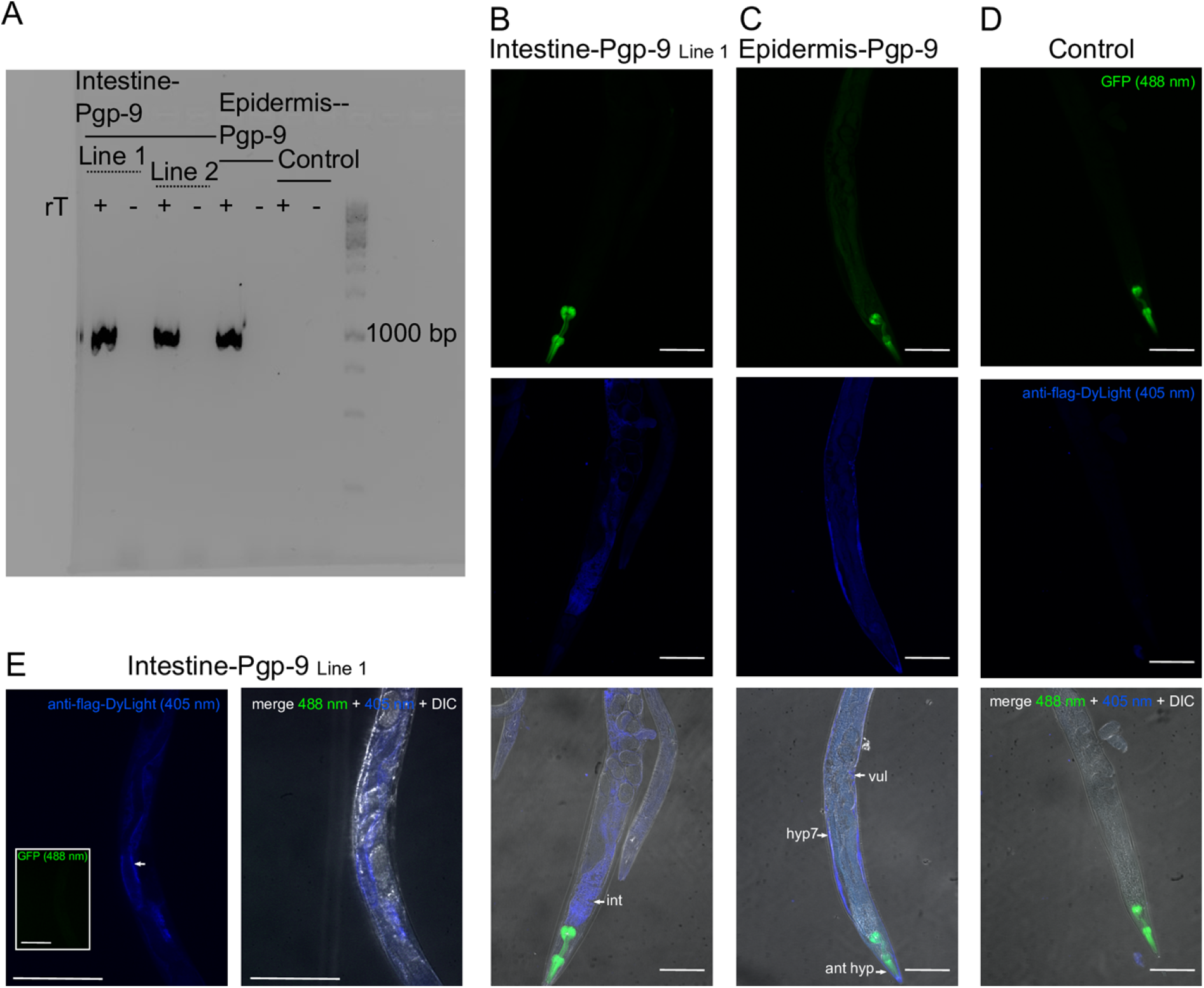
Tissue specific expression of *Pun*-PGP-9 in *Cel-pgp-9* loss of function strain strain tm830. Verification of transcription and tissue-specific expression. (**A**) Visible bands for all reverse-transcriptase PCRs on cDNA templates made from whole worm total RNA and no bands for no RT controls and the control strain. (**B-E**) Confocal microscope images of tissue specific expression patterns of fixed, freeze cracked and immunofluorescence stained transgenic adult *Caenorhabditis elegans* (blue LUT, 405 nM excitation), pharyngeal GFP expression (green LUT, 488 nm excitation) and merged images (DIC +405 nM + 488 nM). Primary antibodies target the FLAG-tag fused to *Pun*-PGP-9 and the secondary antibody is conjugated with DyLight405. Images were acquired with a confocal Eclipse Ti-U inverted research microscope and processed and merged with ImageJ (70). All scalebars are 100 μm. (**B**) Immunostaining in the intestine-Pgp-9 line 1 (white arrows indicate the intestine) and GFP expression at the pharynx. (**C**) Immunostaining of the epidermis-Pgp-9 strain (white arrows indicate the epidermal syncytium) and pharyngeal GFP expression. (**D**) Immunostaining in the control strain and pharyngeal GFP expression. (**E**) Higher magnification of the intestine-Pgp-9 strain (white arrow indicates the apical membrane), *Pun*-PGP-9: *Parascaris univalens* P-glycoprotein-9, GFP: green fluorescence protein, DIC: differential interference contrast, int: intestine, vul: vulva, hyp7: hyp7 syncytium, N2Δ*Cel*Pgp-*9* is tm830 (NBRP) [*Cel-pgp-9(-)*]; Transgenic strains genotypes: Epidermis-Pgp-9 *EpiPgp-9Ex1* [*Cel-pgp-9(-); Cel-col-19p::Pun-pgp-9::FLAG::Cel-unc-54_3′-UTR; Cel-myo-2p::gfp:: Cel-unc-54_3’UTR*]; Intestine-Pgp-9 Line 1 *IntPgp-9Ex1* [*Cel-pgp-9(-); Cel-ges-1p::Pun-pgp-9::FLAG::Celunc-54_3′-UTR; Cel-myo-2p::gfp::Cel-unc-54_3’UTR*]; Control strain *CtrlEx1* [*Cel-pgp-9(-); Cel-myo-2p::gfp:: Cel-unc-54_3′-UTR*]

### 2.2 Pharyngeal pumping differently influences ivermectin and moxidectin susceptibility

Motility assays were conducted as described elsewhere (44) with modifications for stimulation of PP as schematically summarised in (**Fig. 2A**). Prior to drug incubations, induction of PP by OP50 or 4 mM 5-HT or absence of PP in the absence of a stimulus was confirmed visually under a stereo microscope. By PP stimulation, MLs dissolved in DMSO within the incubation medium were actively ingested. After an 18 - 24 h drug incubation, the motility response of worms incubated without drug in 1% DMSO (negative, vehicle control) did not differ between strains or conditions (Kruskal-Wallis test with Dunn’s post-hoc test) compared to the wildtype (WT, untransformed N2 Bristol strain worms) OP50^−^condition (no PP stimulation) (**Fig. 2B**). Therefore, neither transgene expression nor induction of pharyngeal pumping by 5-HT or OP50 significantly influenced motility.

**Figure 2.**
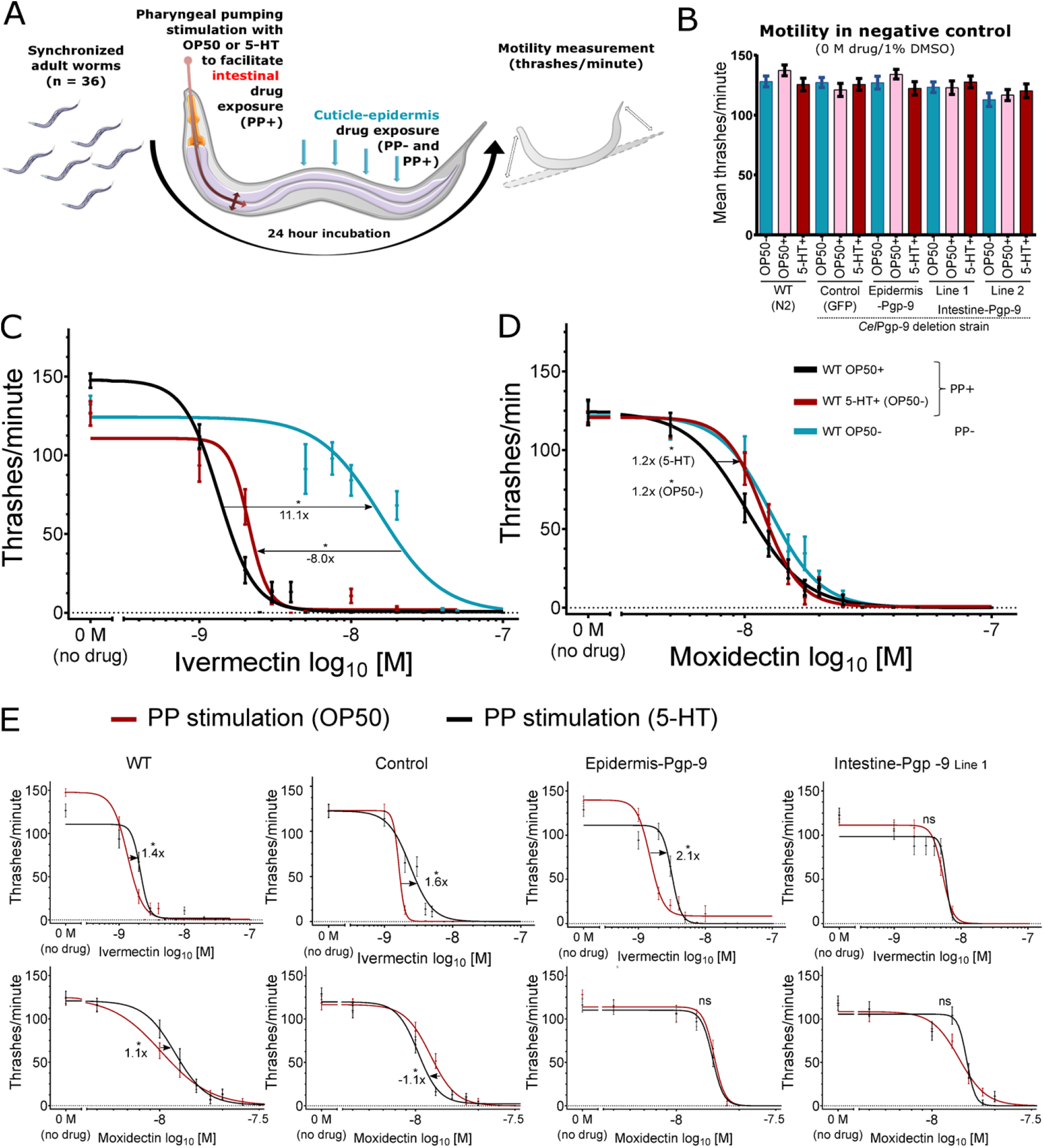
Pharyngeal pumping increases ivermectin and moxidectin susceptibility. Effect of pharyngeal pumping (PP) stimulation by OP50 food bacteria or serotonin in *Caenorhabditis elegans* on ivermectin and moxidectin susceptibility. (**A**) Schematic illustration of the experimental set-up with active ingestion and intestine drug exposure by PP stimulation. (**B**) Mean thrashes/minute ± standard error of the mean (SEM) in the negative control (no drug, 1% DMSO) between strains and conditions. Each strain/condition combination was compared to WT OP50^−^ with a Kruskal-Wallis test with Dunn’s Post-Hoc and p > 0.05 was considered not significant (**C**-**D**) Ivermectin and Moxidectin concentration-response curves calculated with GraphPad v8.3.0 based on thrashes/minute in the WT strain with n = 36 per concentration spread equally on three separate days. Calculated four parameter non-linear regression models (GraphPad v8.3.0, www.graphpad.com) and SEM at each concentration correspond to **Table 1 and 2**. Prior to the calculation, the no drug negative control was set to 0.1 nM and all concentrations were log_10_ transformed. On the x-axis, the negative control was visualized as “0 M (no drug)” and separated by a break in the axis. PP stimulation by OP50 bacteria (OP50^+^) (black), 4 mM 5-HT (red), or in the absence of a PP stimulus (OP50^−^) (blue). Significant differences between EC_50_ were calculated using the extra-sum-of-squares-F test and Bonferroni correction. P-values < 0.05 were considered significant and are indicated with an asterisk while corresponding fold-changes are indicated with arrows. 5-HT: serotonin/5-hydroxytryptamine; WT: *C. elegans* N2 Bristol; Control strain genotype *CtrlEx1* [*Cel-pgp-9(-); Cel-myo-2p::gfp:: Cel-unc-54_3′-UTR*]

Strikingly, the EC_50_ for the WT in the OP50^+^ condition was 1.50 nM IVM, but in the OP50^−^ condition the EC_50_ was 16.68 nM IVM (**Table 1**), which is a significant 11.1-fold increase (p = 0.0014) (**Fig. 2C**). In order to evaluate whether this effect is the result of IVM metabolization by OP50 bacteria or starvation, OP50 were substituted by 4 mM 5-HT in the incubation medium, thereby eliminating the bacteria and food source but maintaining a stimulus for pharyngeal pumping. This resulted in an EC_50_ of 2.08 nM IVM, which is a significant 8.0-fold decrease compared to OP50^−^ (**Fig. 2C**).

**Table 1.**
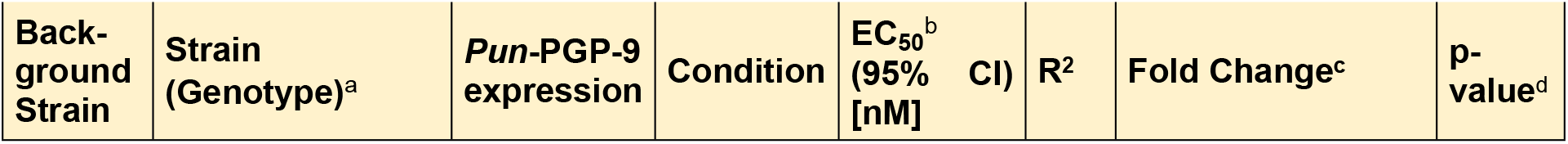
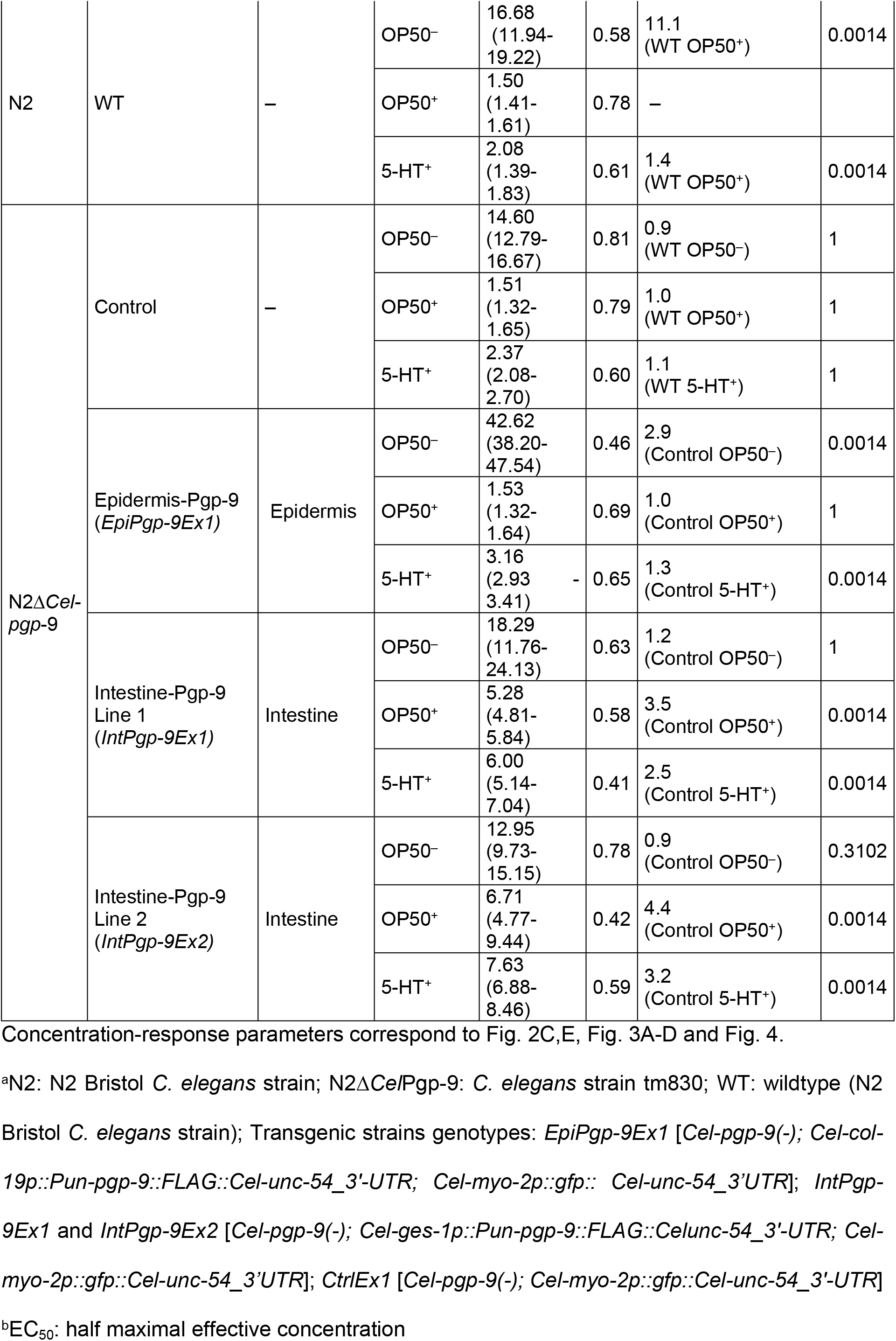
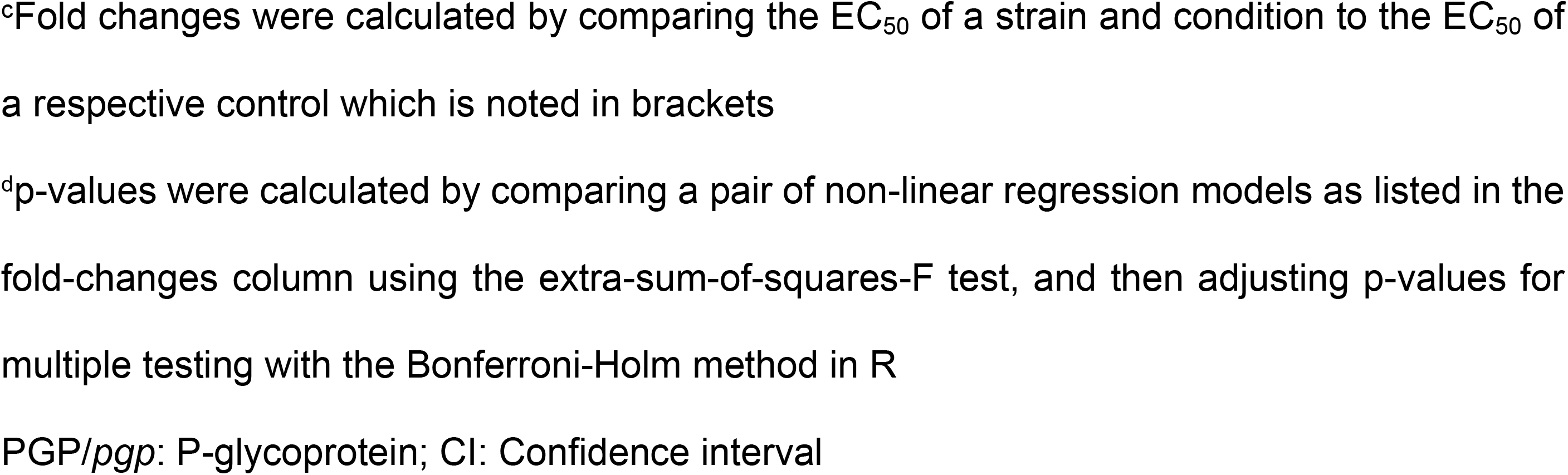
Ivermectin concentration-response parameters of transgenic, control and wildtype strains at different conditions.

| Back-ground Strain | Strain (Genotype)<sup>a</sup> | *Pun*-PGP-9 expression | Condition | $EC_{50}^b$ (95% CI) [nM] | R<sup>2</sup> | Fold Change<sup>c</sup> | p-value<sup>d</sup> |
| --- | --- | --- | --- | --- | --- | --- | --- |
| N2 | WT | – | OP50<sup>-</sup> | 16.68<br>(11.94-<br>19.22) | 0.58 | 11.1<br>(WT OP50<sup>+</sup>) | 0.0014 |
|  |  |  | OP50<sup>+</sup> | 1.50<br>(1.41-<br>1.61) | 0.78 | – |  |
|  |  |  | 5-HT<sup>+</sup> | 2.08<br>(1.39-<br>1.83) | 0.61 | 1.4<br>(WT OP50<sup>+</sup>) | 0.0014 |
| N2Δ*Cel-pgp-9* | Control | – | OP50<sup>-</sup> | 14.60<br>(12.79-16.67) | 0.81 | 0.9<br>(WT OP50<sup>-</sup>) | 1 |
|  |  |  | OP50<sup>+</sup> | 1.51<br>(1.32-<br>1.65) | 0.79 | 1.0<br>(WT OP50<sup>+</sup>) | 1 |
|  |  |  | 5-HT<sup>+</sup> | 2.37<br>(2.08-<br>2.70) | 0.60 | 1.1<br>(WT 5-HT<sup>+</sup>) | 1 |
|  | Epidermis-Pgp-9<br>(*EpiPgp-9Ex1*) | Epidermis | OP50<sup>-</sup> | 42.62<br>(38.20-47.54) | 0.46 | 2.9<br>(Control OP50<sup>-</sup>) | 0.0014 |
|  |  |  | OP50<sup>+</sup> | 1.53<br>(1.32-<br>1.64) | 0.69 | 1.0<br>(Control OP50<sup>+</sup>) | 1 |
|  |  |  | 5-HT<sup>+</sup> | 3.16<br>(2.93-<br>3.41) | 0.65 | 1.3<br>(Control 5-HT<sup>+</sup>) | 0.0014 |
|  | Intestine-Pgp-9<br>Line 1<br>(*IntPgp-9Ex1*) | Intestine | OP50<sup>-</sup> | 18.29<br>(11.76-24.13) | 0.63 | 1.2<br>(Control OP50<sup>-</sup>) | 1 |
|  |  |  | OP50<sup>+</sup> | 5.28<br>(4.81-<br>5.84) | 0.58 | 3.5<br>(Control OP50<sup>+</sup>) | 0.0014 |
|  |  |  | 5-HT<sup>+</sup> | 6.00<br>(5.14-<br>7.04) | 0.41 | 2.5<br>(Control 5-HT<sup>+</sup>) | 0.0014 |
|  | Intestine-Pgp-9<br>Line 2<br>(*IntPgp-9Ex2*) | Intestine | OP50<sup>-</sup> | 12.95<br>(9.73-15.15) | 0.78 | 0.9<br>(Control OP50<sup>-</sup>) | 0.3102 |
|  |  |  | OP50<sup>+</sup> | 6.71<br>(4.77-<br>9.44) | 0.42 | 4.4<br>(Control OP50<sup>+</sup>) | 0.0014 |
|  |  |  | 5-HT<sup>+</sup> | 7.63<br>(6.88-<br>8.46) | 0.59 | 3.2<br>(Control 5-HT<sup>+</sup>) | 0.0014 |

Concentration-response parameters correspond to Fig. 2C,E, Fig. 3A-D and Fig. 4.
<sup>a</sup>N2: N2 Bristol *C. elegans* strain; N2Δ*CelPgp-9*: *C. elegans* strain tm830; WT: wildtype (N2 Bristol *C. elegans* strain); Transgenic strains genotypes: *EpiPgp-9Ex1* [*Cel-pgp-9*(-); *Cel-col-19p::Pun-pgp-9::FLAG::Cel-unc-54\_3'-UTR*; *Cel-myo-2p::gfp:: Cel-unc-54\_3'UTR*]; *IntPgp-9Ex1* and *IntPgp-9Ex2* [*Cel-pgp-9*(-); *Cel-ges-1p::Pun-pgp-9::FLAG::Celunc-54\_3'-UTR*; *Cel-myo-2p::gfp::Cel-unc-54\_3'UTR*]; *CtrlEx1* [*Cel-pgp-9*(-); *Cel-myo-2p::gfp::Cel-unc-54\_3'-UTR*]
<sup>b</sup>EC<sub>50</sub>: half maximal effective concentration
<sup>c</sup>Fold changes were calculated by comparing the EC<sub>50</sub> of a strain and condition to the EC<sub>50</sub> of a respective control which is noted in brackets
<sup>d</sup>p-values were calculated by comparing a pair of non-linear regression models as listed in the fold-changes column using the extra-sum-of-squares-F test, and then adjusting p-values for multiple testing with the Bonferroni-Holm method in R
PGP/*pgp*: P-glycoprotein; CI: Confidence interval

For MOX, stimulation of PP by OP50 also resulted in a significantly lower EC_50_ than in the absence of PP (OP50^−^). Specifically, EC_50_ values in the WT strain in the presence of PP stimulation were 10.19 nM MOX (OP50^+^) and 11.85 nM MOX (5-HT^+^) (**Table 2**). However, this change was only by 1.2-fold and thus considerably smaller than the effect of PP on IVM susceptibility. PP stimulation by 5-HT did not significantly increase susceptibility, and the EC_50_ was by 1.2-fold significantly higher than stimulation by OP50.

**Table 2.**
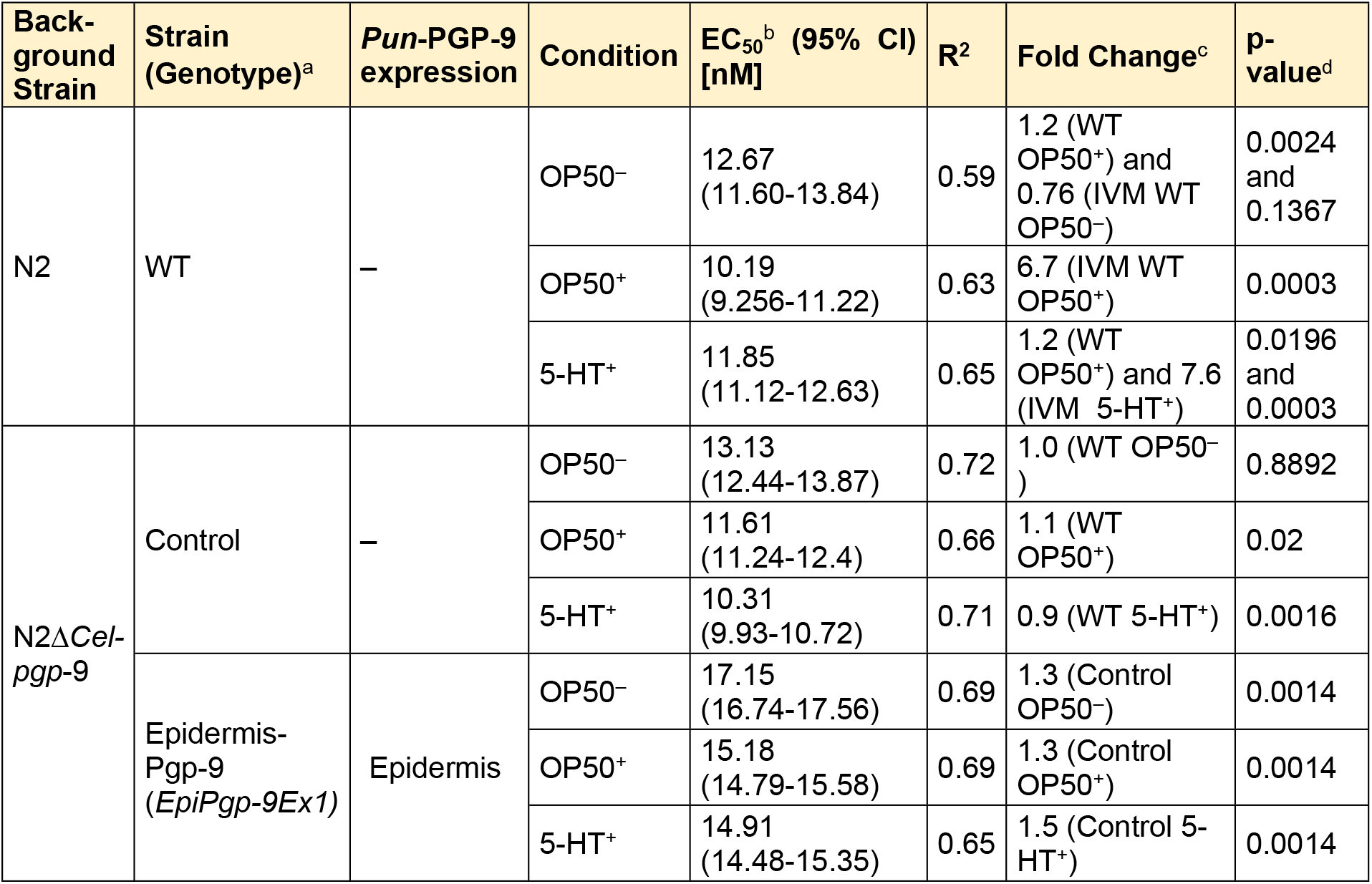
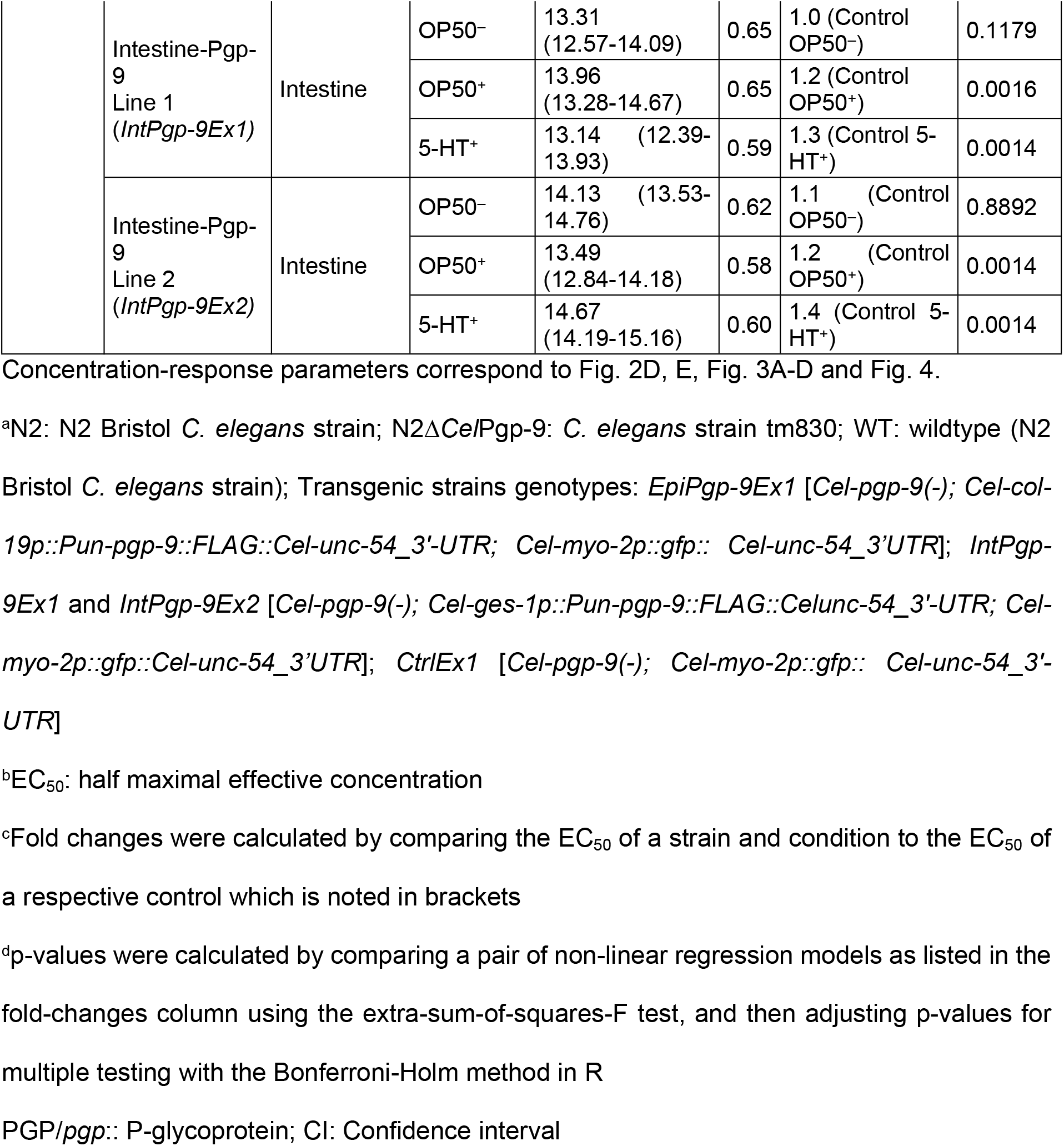
Moxidectin concentration-response parameters of transgenic, control and wildtype strains at different conditions.

Across all strains except intestine-Pgp-9, IVM EC_50_ values for PP induction by 5-HT were higher compared to PP stimulation by OP50 (**Fig. 2E**) (p < 0.05, extra-sum-of-squares F test, Bonferroni corrected). In contrast, between 5-HT and OP50 PP stimulation MOX EC_50_ values varied slightly in the control (*Cel-pgp-9* null mutant with pharyngeal GFP expression) and WT, and these strains exhibited a 1.1-fold higher (WT) or a 1.1-fold lower (control) EC_50_ value (p < 0.05, extra-sum-of-squares F test, Bonferroni corrected, 5-HT^+^ compared to OP50^+^) while no significant difference was present in the epidermis-Pgp-9 and intestine-Pgp-9 strains.

### 2.3 The adult stage motility phenotype in response to ivermectin or moxidectin is unaffected by *Cel-pgp-9* loss-of-function

The control strain which is *Cel-pgp-9* null mutant (*CtrlEx1 [Cel-pgp-9(-); Cel-myo-2p::gfp:: Cel-unc-54_3′-UTR*]) did not exhibit increased susceptibility to IVM or MOX (**Supplementary Fig. 1A and B**). Serving as a control for extrachromosomal array transgene expression (pharyngeal GFP expression), the calculated EC_50_ values in the control strain matched those of the WT for IVM (p = 1, **Table 1**). Likewise, the control strain’s response to MOX was similar to that of the WT strain, and despite minor variation between the two strains, MOX EC_50_ values were all within a close range (**Table 2**). Generally, PP stimulation slightly increased MOX susceptibility (**Table 2**). The control strain was used as a reference for both the *Pun*-PGP −9 overexpression strains, epidermis-Pgp-9 and intestine-Pgp-9, as the control strain exhibits the same genetic background (*Cel-pgp-9* loss of function) and transgenic extrachromosomal expression.

### 2.4 Epidermal *Pun*-PGP-9 expression reduces ivermectin and moxidectin susceptibility

Concerning IVM susceptibility, the epidermis-Pgp-9 line resembled the control in the presence of an OP50 associated PP stimulus (p = 1) (**Table 1**) while in the presence of a 5-HT PP stimulus the EC_50_ was by 1.3-fold slightly higher compared to incubation with OP50 (**Fig. 3A**, p = 0.0014). Strikingly, in the absence of a PP stimulus, the EC_50_ increased significantly to 42.62 nM IVM, which is 2.9-fold higher than the control strain for the same conditions (p = 0.0014).

**Figure 3.**
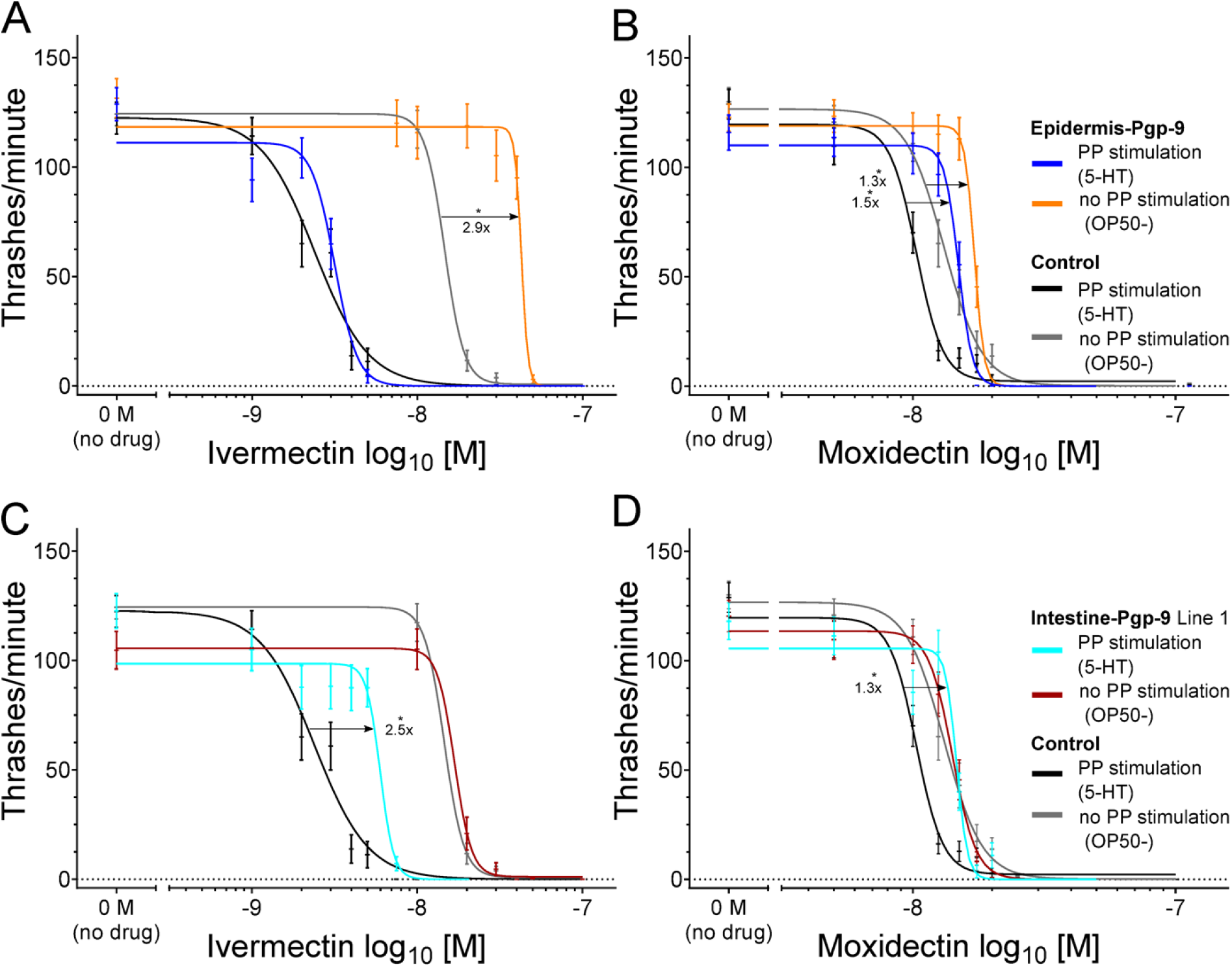
Modulation of ivermectin and moxidectin susceptibility in C. elegans by transgenic tissue-specific Pun-PGP-9 expression. (**A**-**D**) Concentration-response curves corresponding to **Table 1** **and** **2** for ivermectin (**A**/**C**) and moxidectin (**B**/**D**) in the absence of a PP stimulus (OP50^−^) or PP stimulation by serotonin (5-HT^+^) in liquid S-medium. (**a**-**D**) show the control strain’s response in black (5-HT^+^) or grey (OP50^−^), (**a**-**b**) show the epidermis-Pgp-9 strain in blue – (5-HT^+^) or orange (OP50^−^) and (**C**-**D**) show the intestine-Pgp-9 line 1 in turquoise (5-HT^+^) or red (OP50^−^). Concentration-response curves are based on four parameter non-linear regression models (GraphPad v8.3.0, www.graphpad.com) calculated from the motility response (thrashes/minute) of 36 synchronized 1-day adults per concentration and error bars represent standard error of the mean. Prior to calculation, concentrations were log_10_ transformed and the no drug negative control was set to 0.1 nM. On the x-axis, the negative control was visualized as “0 M (no drug)” and separated by a break in the axis. Significant differences in EC_50_ values were compared using the extra-sum-of-squares-F test and Bonferroni correction p-values < 0.05 were considered significant and indicated with an asterisk (*) while corresponding fold-changes are indicated with arrows. *Pun*-PGP-9: *Parascaris univalens* P-glycoprotein-9; N2Δ*Cel*Pgp-*9* is tm830 (NBRP) [*Cel-pgp-9(-)*]; Transgenic strains genotypes: Epidermis-Pgp-9 *EpiPgp-9Ex1* [*Cel-pgp-9(-); Cel-col-19p::Pun-pgp-9::FLAG::Cel-unc-54_3′-UTR; Cel-myo-2p::gfp:: Cel-unc-54_3’UTR*]; Intestine-Pgp-9 Line 1 *IntPgp-9Ex1* [*Cel-pgp-9(-); Cel-ges-1p::Pun-pgp-9::FLAG::Celunc-54_3′-UTR; Cel-myo-2p::gfp::Cel-unc-54_3’UTR*]; Control strain *CtrlEx1* [*Cel-pgp-9(-); Cel-myo-2p::gfp:: Cel-unc-54_3′-UTR*]

Epidermal *Pun*-PGP-9 expression also impacted MOX susceptibility (**Table 2**). Similarly to IVM, in the absence of PP the epidermis-Pgp-9 strain exhibited the highest EC_50_ value (17.15 nM MOX for OP50^−^) across strains and conditions (**Table 2**). Interestingly, epidermal *Pun*-PGP-9 expression also resulted in the overall lowest MOX susceptibility between all strains regardless of PP stimulation with EC_50_ values of 15.18 nM MOX (OP50^+^) and 14.91 nM MOX (5-HT^+^) (**Table 2, Fig. 3B**). Notably, the differences to the control strain are only moderate, i.e. a 1.3-fold in the absence of PP (OP50^−^) and a 1.3-fold (OP50^+^) or 1.5-fold (5-HT^+^) increase in the presence of PP stimulation.

In summary, epidermal *Pun*-PGP-9 expression reduced susceptibility to IVM only in the absence of PP stimulation but to MOX regardless of PP.

### 2.5 Reduction of ivermectin and moxidectin susceptibility from intestinal *Pun*-PGP-9 expression only in the presence of pharyngeal pumping

Remarkably, intestinal *Pun*-PGP-9 expression reduced the susceptibility to IVM in the presence of a PP stimulus compared to the control strain (**Fig. 3C**) in two independent lines (**Supplementary Fig. 1 C and D)**. In detail, in the presence of OP50 bacteria or 5-HT PP stimulation, the EC_50_ values were significantly increased by 3.5- and 2.5-fold in line 1 and 4.4- and 3.2-fold in line 2 (all p = 0.0014) (**Table 1**), compared to the control strain. In contrast, EC_50_ values for both lines were not significantly different from the control strain when PP was not stimulated (p = 1; p = 0.3102) (**Table 1** **and** **Fig. 3C**).

MOX EC_50_ values in the two intestine-Pgp-9 lines were also significantly elevated when PP was stimulated by OP50 (p = 0.0016 and 0.0014) or 5-HT (both p = 0.0014) compared to the control strain (**Table 2** **and** **Fig. 3D**). However, in contrast to IVM, the corresponding fold-changes were considerably smaller (<1.3-fold **Table 2**). In addition, all MOX EC_50_ values for the two lines were overall lower than the MOX EC_50_ values calculated for the epidermis-Pgp-9 line. In the absence of PP stimulation, EC_50_ values did not increase compared to the control strain, which is in line with the observations for IVM.

In summary, intestinal *Pun*-PGP-9 expression reduced susceptibility to both tested MLs but this effect was always dependent on active drug ingestion.

### 2.6 A comparison of moxidectin and ivermectin susceptibility in transgenic and wild-type strains

Compared to IVM, EC_50_ values for MOX were generally higher across strains when PP was stimulated but lower in the absence of a PP stimulus (**Fig. 4**) as detailed below. For example, MOX EC_50_ values in the WT in the presence of PP stimulation were significantly higher, by 6.7- and 7.6-fold (both p = 0.0003), compared to IVM (**Table 2** **and** **Fig. 4**) for co-incubation with OP50 or 5-HT, respectively. In contrast, when a PP stimulus was not provided, the MOX EC_50_ was not significantly different compared to IVM (p = 0.1367) (**Table 2**, **Fig. 4**). Across strains, only small fold-changes below 1.2-fold for the MOX EC_50_ values were detected in the presence or absence of PP stimulation (**Table 2**), which is in a sharp contrast to IVM. As for IVM, a marked EC_50_ increase (up to 11-fold in WT) between PP and no PP was observed in all strains (**Fig. 4**). Likewise, *Pun*-PGP-9 expression did only result in low fold changes for MOX EC_50_ values (<1.5-fold) in comparison to considerably higher fold changes (e.g. >3.5-fold in intestine-Pgp-9) for IVM (**Table 1** **and** **Fig. 4**). Despite the marked effect on IVM susceptibility from intestinal *Pun*-PGP-9 overexpression, all EC_50_ values for both intestine-Pgp-9 lines in the presence of PP stimulation remained lower compared to EC_50_ values of any other strain in the absence of a PP stimulus (**Fig. 4**). In contrast, MOX EC_50_ values in the intestine-Pgp-9 strains in the presence and absence of PP were similar (**Fig. 4**).

**Figure 4.**
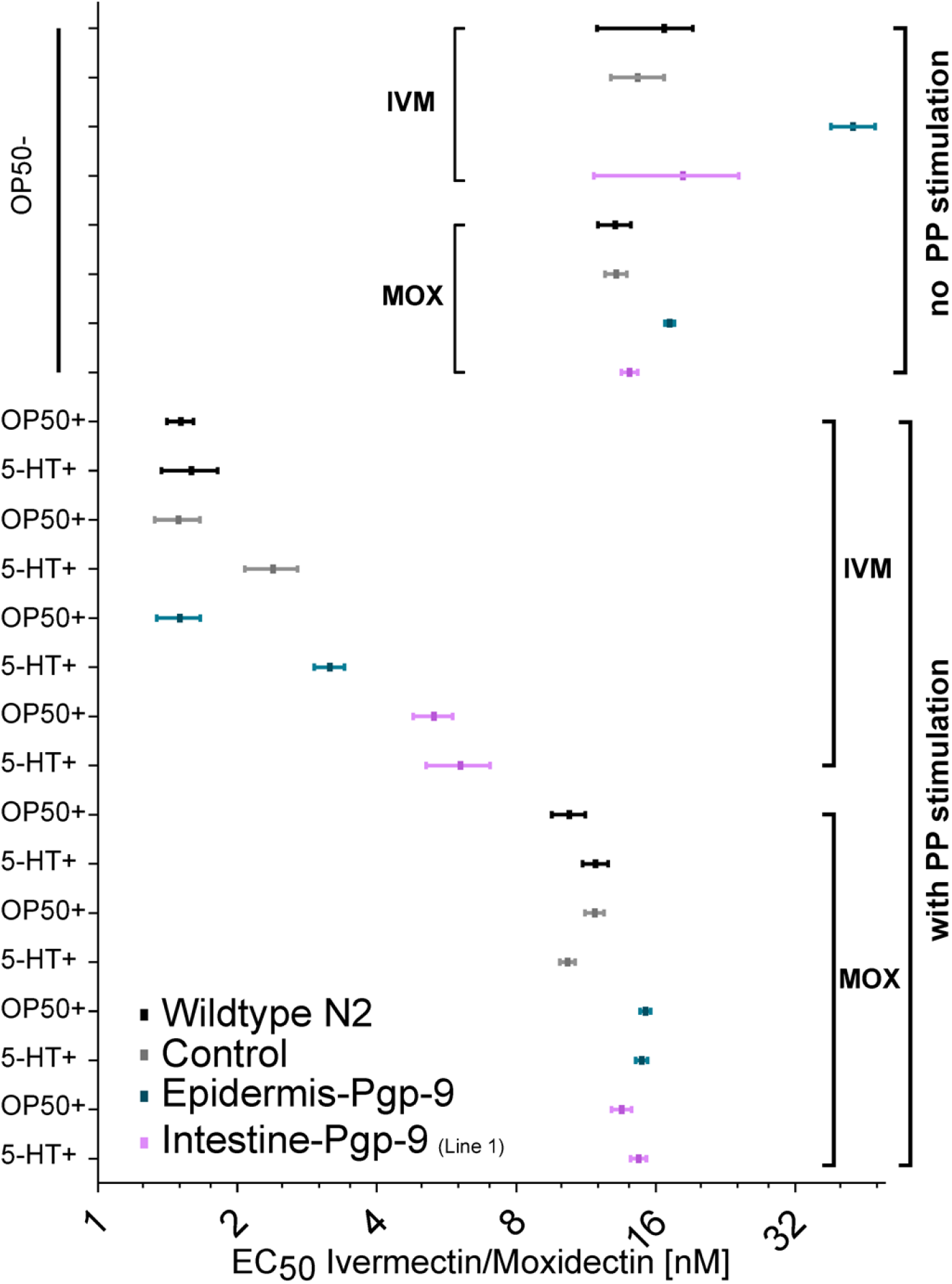
Comparison of moxidectin and ivermectin in wildtype and transgenic *Caenorhabditis elegans* strains. Forrest plot visualising the EC_50_ and corresponding 95% confidence intervals in nM on a log_2_ axis for both ivermectin and moxidectin in transgenic and wildtype *Caenorhabditis elegans* strains in the presence of PP stimulation by OP50 food bacteria (OP50^+^) or serotonin (5-HT^+^) or no PP stimulation (OP50^−^). EC_50_ values were inferred from four parameter linear regression models calculated from 36 synchronized 1-day adult worms per concentration, strain and condition. Worms were incubated for 24 hours in S-medium containing a concentration series of ivermectin or moxidectin in a final DMSO concentration of 1%. Epidermis-Pgp-9 (turquoise) genotype is *EpiPgp-9Ex1* [*Cel-pgp-9(-); Cel-col-19p::Pun-pgp-9::FLAG::Cel-unc-54_3′-UTR; Cel-myo-2p::gfp:: Cel-unc-54_3’UTR*];. Intestine Pgp-9 line 1 (purple) genotype is *IntPgp-9Ex1* [*Cel-pgp-9(-); Cel-ges-1p::Pun-pgp-9::FLAG::Celunc-54_3′-UTR; Cel-myo-2p::gfp::Cel-unc-54_3’UTR*];. Control strain (grey) genotype is Control strain *CtrlEx1* [*Cel-pgp-9(-); Cel-myo-2p::gfp:: Cel-unc-54_3′-UTR*]; WT (N2 Bristol) (black). Pgp: P-glycoprotein; *Pun*-PGP-9: *Parascaris univalens* P-glycoprotein-9; IVM: Ivermectin; MOX: Moxidectin; PP: Pharyngeal pumping. WT: wildtype.

## 3 Discussion

Active drug ingestion by PP markedly increased IVM and, to a lesser degree, MOX susceptibility in *C. elegans*, suggesting that PP increases the overall ML uptake. In this context, the role of tissue specific Pgp expression was investigated, revealing novel insights into the mechanism of Pgp-mediated ML resistance. In fact, this is the first direct functional evidence of a transgenic, parasitic nematode derived Pgp causing reduced IVM and MOX susceptibility in a tissue-specific manner. The dependency of this protective effect on PP suggests a role of Pgp in barrier fortification as a possible mechanism of a Pgp-mediated ML susceptibility. Concurrently, the different susceptibilities between MOX and IVM indicate differences in their affinity to *Pun*-PGP-9 and, importantly, in their ability to penetrate the intestinal and the cuticular-epidermal barrier.

While Pgp have been linked to ML resistance in numerous studies, the functional mechanisms by which they contribute to the ML resistance in nematodes remained unknown (16, 45). Here, we provide a direct link between Pgp overexpression in a nematode and reduced ML susceptibility and shed light on the functional role of tissue-specific Pgp expression. Our findings support the hypothesis that Pgps contribute to barrier function at the intestine and the cuticle-epidermis as illustrated in **Fig. 5**. In contrast, if excretion via the excretory system were the relevant mechanism, a dependency on PP would not be expected for the protective effect of drug efflux by overexpressed transgenic Pgps. Noteworthy, intestinal excretion is dependent on PP as defecation is halted in the absence of PP. Our conceptual discrimination between these two mechanisms is that excretion would occur regardless of the penetration route, e.g. intestinal excretion following cuticular-epidermal drug penetration, while barrier function is the prevention of drug penetration directly at the uptake tissue. In case of excretion, regardless of the drug entry route, increased elimination by *Pun*-PGP-9 via the intestine or the epidermal excretory pore should elicit a protective effect. For IVM the strong impact of PP on susceptibility and the complete dependency of the effect of tissue-specific Pgp expression point towards a role in barrier function. These mechanistic hypotheses for Pgp function in nematodes are inspired by the role of Pgps in mammals. Here, Pgps stop MLs from crossing the blood-brain barrier, and at the intestine, they contribute to ML elimination by excretion (27, 28). In nematodes, Pgp expression is predominant in the intestine (32, 35, 37) suggesting a role in excretion or barrier function in this tissue. Pgp expression in the epidermis has also been reported in several nematode species at a moderate level e.g., in *Parascaris* spp. (35), and *C. elegans* (46) but their role in this tissue has not received much attention. Notably, Pgps are also expressed in the excretory system, particularly the H-shaped cell, as well as in the neuronal system in both *C. elegans* (46) and *Parascaris* spp. (35). Whether Pgp expression in these tissues by ML excretion via the H-shaped excretory cell, the intestine, and the epidermis, or even Pgp co-localization with ML targets in the membranes of nerve cells elicits protective effects remains to be elucidated.

**Figure 5.**
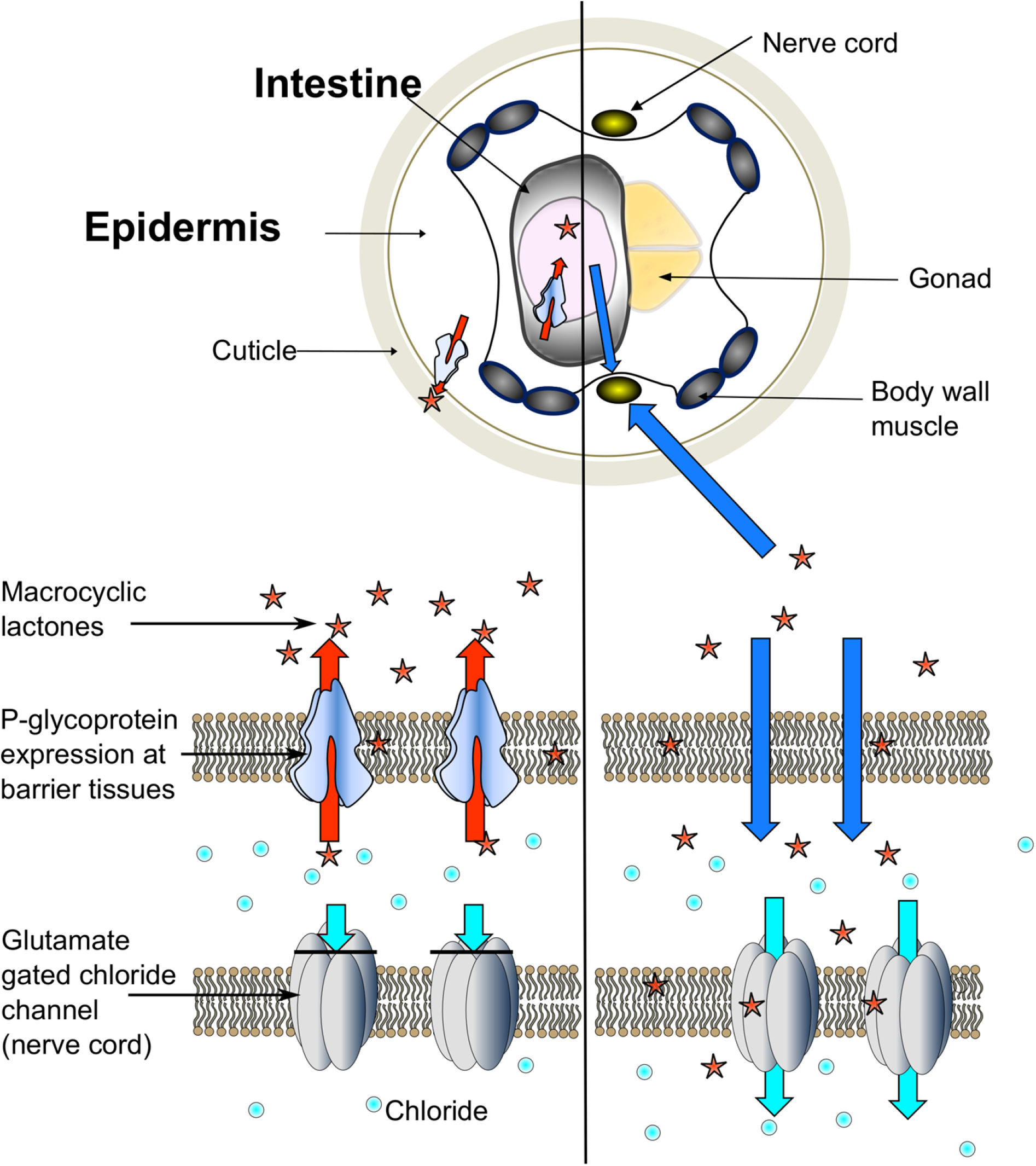
Schematic illustration of P-glycoprotein-mediated barrier function. Hypothetical schematic illustration of Pgp-mediated barrier function in a *Caenorhabditis elegans* adults. Expression of P-glycoproteins in specific barrier tissues, i.e. the epidermis and the intestine prohibit MLs from reaching target tissues thereby preventing an ML induced hyperpolarisation of the neurons and muscle paralysis.

An important question addressed in this work was whether there are differences in the protective effect of Pgp overexpression between different ML derivatives. Currently, functional evidence regarding the interaction between parasite Pgps and MLs has mostly been inferred from indirect assays, which use MLs as inhibitors for a secondary Pgp substrate with a discernible effect in the model systems to characterize the interaction between a transgenic parasite Pgp and different MLs. Thus, these approaches assess the Pgp ML interaction relative to the secondary substrate’s affinity to the Pgps, for example, the antimycotic ketoconazole in transgenic yeast or the fluorescent dye rhodamine 123 in the cell line LLC-PK1 (25, 37). Through these indirect binding assays several nematode parasite Pgps such as *Cylicocyclus elongatus* PGP-9 (25), *Pun-*PGP −2, and *Pun*-PGP-9 (32), *D. immitis* PGP-11 (47) and *H. contortus* PGP-2, −9.1 or −16 (24, 37, 48) have been characterized. These studies have consistently suggested that MOX has a lower affinity to nematode Pgps than IVM. This conclusion is further supported by our findings in *C. elegans* that *Pun*-PGP-9 overexpression induces larger susceptibility shifts for IVM than MOX which were observed without the use of a secondary substrate. However, *Pun*-PGP-9 overexpression reduced both MOX and IVM susceptibility, which indicates that both MLs are *Pun*-PGP-9 substrates. For nematode populations where Pgp-mediated ML resistance represents an important component of their resistance mechanism, cross-resistance between the two derivatives might thus occur. Indeed, resistance against multiple MLs is common in *Parascaris* spp. (49). In *C. elegans*, only one study had previously demonstrated the impact of a transgenic Pgp on IVM susceptibility, i.e., *Pun-*PGP-11.1 expressed under the control of the native *Cel-pgp-11* promotor (44). The reported reduction in IVM susceptibility by 4.6- and 3.2-fold in two separate lines was similar to the observed shifts in this study in the intestine-Pgp-9. In addition, the authors used similar incubation conditions and always included OP50 in their incubation media (44). As the *Cel-pgp-11* promotor has been shown to primarily drive intestinal expression (50), the findings of Janssen et al. are in high concordance with the observations made in this study.

This study is the first to identify differences in the “pharmacokinetics” of MLs in nematodes. The results reveal the effects of active drug ingestion and the PP dependent effect of tissue-specific transgenic Pgp overexpression on susceptibility. These differences point towards differing IVM and MOX uptake capacities at specific *C. elegans* tissues, which to the authors’ best knowledge is unprecedented. As a general mechanism, PP stimulation is required for food uptake, and the increased uptake of incubation fluid also facilitates the accumulation of its contents (40). For IVM, the strong PP dependency on the protective effect of *Pun*-PGP-9 indicates that higher intestinal exposure increases the overall dose of IVM. Although the uptake capacities cannot be directly quantified, a possible interpretation of the combined effects of PP and tissue-specific *Pun*-PGP-9 expression is that IVM is taken up more efficiently via the intestine. Specifically, this is supported by the strong decrease in IVM susceptibility in the absence of a PP stimulus observed in all of the examined *C. elegans* strains. In line with this conclusion, intestinal *Pun*-PGP-9 overexpression resulted in a marked decrease in IVM susceptibility in two independent lines in the presence of PP stimulation. Concurrently, the impact of epidermal *Pun*-PGP-9 expression on susceptibility and the lack of an effect of intestinal expression on susceptibility in the absence of PP also indicate that IVM is taken up, although less efficiently, via the cuticle-epidermis. It appears that this is the relevant uptake route when PP is not stimulated. In contrast, fold shifts in MOX susceptibilit between strains and PP stimulation differ from IVM. Here, the effect of epidermal *Pun*-PGP-9 expression on MOX susceptibility regardless of PP could suggest that the epidermis-cuticle is the predominant uptake route.

For IVM and other anthelmintics, trans-cuticular-epidermal permeability has also been suggested by biophysical studies on cuticle-epidermis preparations of *Ascaris suum* and experiments using live *A. suum* with surgically ligated pharynx, but PP was not stimulated in the unligated worms. Thereby, the authors might have underestimated the importance of active drug ingestion and intestinal uptake (51, 52). Likewise, experiments in *C. elegans* have demonstrated a considerably quicker onset of pharyngeal paralysis in a cuticle defective *C. elegans bus-8* loss-of-function strain (53). In line with the findings of this study, a study by O’Lone and Campbell indicated that PP inhibition by refrigeration and thus inhibition of oral ingestion reduced the susceptibility to IVM in *C. elegans* (54). The authors suggested that oral ingestion increases IVM susceptibility but that transcuticular-epidermal uptake is sufficient to induce paralysis at comparatively low concentrations (54). These conclusions are in high concordance with our observations and conclusions.

Conversely, we also demonstrate the direct effect of active PP stimulation on IVM susceptibility. This effect is considerably smaller for MOX, highlighting the need for a better understanding of drug penetration through specific nematode tissues and the potential differences between different compounds. The uptake of MOX in nematodes has not been studied so far, but a possible interpretation of the results in the present study is that the cuticle-epidermis represents the main drug penetration route. This conclusion would be supported by the higher MOX concentrations than IVM needed to induce paralysis when PP was stimulated. Similarly, another study using WT *C. elegans* reported a lower susceptibility to MOX compared to IVM for motility and other read-out phenotypes (55). In contrast, when active oral ingestion was not induced, MOX induced paralysis occurred at lower concentrations than IVM. In addition, the epidermal transgenic Pgp overexpression resulted in the overall lowest MOX susceptibility regardless of PP stimulation. Concurrently, the PP dependent effect of intestinal transgenic Pgp overexpression suggests that the drug can also be taken up via the intestine.

These differences must be the result of the differing chemico-physical properties of IVM and MOX. The main factor determining permeability through the nematode cuticle is compound size (51, 52, 56). and MOX is considerably smaller than IVM due to the lack of the C13 glycosyl side chain. Additionally, lipophilicity is an ancillary but important factor for transcuticular permeability (51, 52, 56), and indeed IVM has a lower lipophilicity than MOX (57). These difference in size and lipophilicity seem to offer a reasonable explanation for a less efficient transcuticular IVM uptake. Both ML derivatives are very lipophilic and accumulate well in high-fat content tissues in the host (58). In this regard, the mainly intestinal and epidermal fat storage in *C. elegans* (59) should facilitate passive diffusion and accumulation in these tissues.

The overall moderate fold changes observed in this study result from the brute force extra-chromosomal overexpression of *Pun*-PGP-9. While expression was not quantified, even strong Pgp overexpression in wild populations is unlikely to completely explain ML resistance in parasitic nematodes. However, these results confirm the potential of Pgps to reduce ML susceptibility and contribute to a protective barrier. It should be noted that the novel insights from the present study are, to some extent, limited to the model nematode *C. elegans*. Parasitic nematodes exhibit differences in size, anatomy, and genetics, e.g., in their receptor repertoire (60) and composition and hence drug susceptibility (61). For example, IVM and MOX EC_50_ values for adult *C. elegans* are several folds higher than those measured in the adult stage of examined parasitic nematodes, e.g., *T. circumcinta* (62). The reasons for these differences remain elusive and have been a matter of speculation. In non-feeding third larval stages, exposure to MLs (or any other anthelmintic) is restricted to the cuticle. and indeed non-feeding larval stages of parasitic nematodes exhibit considerably lower susceptibilities (62, 63). This is in line with the observed increase in ML susceptibility from PP in this study. Furthermore, the sheath (i.e., the residual cuticle of the second larval stage) of the third stage strongylid larva might also reduce susceptibility by limiting drug penetration into the worm. Regarding the uptake of MLs in adult stages of parasitic worms, an *in vivo* study on ML resistant *H. contortus* found no difference of initial drug accumulation in parasites between MOX, IVM, and abamectin 0.5 days after treatment (64). However, MOX concentrations dropped significantly compared to IVM and abamectin within two days post treatment, suggesting that drug elimination may vary in parasitic nematodes between different ML derivatives. Importantly, the drug pharmacokinetics in the host also varied substantially between different ML derivatives and did not correlate with drug levels in the parasites, highlighting the knowledge gap between drug pharmaco-kinetics and -dynamics in the host and in the parasite. Exposure duration and dose considerably differ between parasitic species and stages, e.g., migrating stages. For intestinal dwelling parasitic nematodes, feeding behaviour would be expected to influence the exposure dose. For example, the blood sucking *H. contortus* would experience high intestinal ML exposure during the early peak in plasma concentrations and thereafter prolonged low intestinal exposure, if considering the pharmacokinetic ML profile of subcutaneous formulations (65). In turn, the cuticle epidermis would only be exposed from the surrounding gastrointestinal fluid and the accumulation in the intestine or abomasal content varies considerably between ML derivatives and drug formulation (64, 66). Our findings emphasize that more attention should be placed on how target parasite species take up ML drugs since this might lead to differences in susceptibility to individual ML derivatives. Whether specific barrier changes represent a relevant mechanism of anthelmintic resistance, could be answered by analyses of tissue specific gene expression changes or histological examination in the context of anthelmintic resistance.

In conclusion, this study significantly improves the understanding of a Pgp-mediated ML resistance mechanism by demonstrating how transgenic Pgp expression at specific barriers can impact the susceptibility to different ML derivatives. Furthermore, the relevance of PP for ML susceptibility *in C. elegans* suggests pharmacological differences in tissue uptake capacities between two important ML derivatives, MOX and IVM.

## 4 Materials and methods

### 4.1 Plasmids and plasmid construction

Briefly, plasmid constructs for transgenesis *Cel*p*col-19:: Pun-pgp-9::FLAG::Celunc-54_3′-UTR* driving epidermal *Pun*-PGP-9 (**Supplementary Fig. 1A**) expression and *Celpges-1:: Pun-pgp-9::FLAG::Celunc-54_3′-UTR* (**Supplementary Fig. 1B**) driving intestinal *Pun*-PGP-9 expression were assembled using the NEB HIFI DNA Assembly Kit (New England Biolabs Inc.) according to the manufacturer’s instructions into the supplied pUC19 vector linearized by *Sma*I (ThermoFisher) digestion. The *C. elegans unc-54* 3′-UTR, the promotor p*ges-1* (intestine) (44) and the *Pun-pgp-9* gene (32) were amplified from a plasmid while the 3′ end primer for the *Pun-pgp-9* amplification introduced an in-frame FLAG-tag (DYKDDDDK) before the stop codon (all Primers in Supplementary Table 1). The *C. elegans* promotor p*col-19* (epidermis) was amplified from genomic DNA extracted from the Bristol N2 strain (NucleoSpin Tissue XS kit (Macherey Nagel) (43). A co-injection marker plasmid (pPD118.33) driving pharyngeal GFP expression was used (Addgene L3790, a gift from A. Fire). Sequences of all constructs were confirmed by Sanger-sequencing (LGC Genomics).

### 4.2 Generation of *Caenorhabditis elegans* strains and maintenance

*Caenorhabditis elegans* WT strain Bristol N2 was obtained from the Caenorhabditis Genetics Center (CGC; University of Minnesota, Minneapolis, MN, USA) and the *Cel-pgp-9* deletion strain tm830 [*Cel-pgp-9(-)*] from the National BioResource Project (NBRP; Tokyo, Japan). Strains were maintained at 20 °C and standard conditions on NGM plates (67).

Plasmid constructs for intestinal and epidermal *Pun*-PGP-9 expression were loaded into Eppendorf FemtoTips II needles at 25.0 ng/μL along with pPD118.33 as a co-injection marker at 12.5 ng/μL. Injection into the gonads of adults of the *Cel-pgp-9* deletion strain (tm830) was performed using an Eppendorf Femtojet 4 connected to a Eppendorf micromanipulator and mounted onto a Leica inverse microscope. Preparation of worms for injection was carried out as described elsewhere (68). Additionally, a strain controlling for pharyngeal GFP expression and presence of extra-chromosomal arrays in general was generated by injecting only the co-injection marker. Transgenic strains were maintained by regular transfer of GFP positive individuals to a new plate.

### 4.3 Verification of *Pun*-PGP-9 expression by RT-PCR and Immunofluorescence

Transcription of *Pun-pgp-9* in GFP positive offspring of injected worms was confirmed by reverse transcriptase (RT)-PCR (primers and cycle conditions in Supplementary Table 1) using the Phusion S7 enzyme (Mobidiag). In addition, tissue specific *Pun*-PGP-9 protein expression was examined with immunofluorescence. Freeze cracking and antibody staining were performed in a 1.5 ml tube similarly as described elsewhere (69) with minor adaptations using 4% formaldehyde and 50% Methanol for fixation and freeze-cracking worms three times in liquid nitrogen. Following three freeze-thaw cycles, tubes were shaken at 37 °C for 1 hour. To remove the fixative, worms were centrifuged at 11,000 × g for 1 minute and washed with PBS-T (phosphate buffered saline + 0.5% Triton-X100) four times, removing all liquid during the last aspiration of PBS-T. Prior to antibody staining, worms were incubated in 1 ml PBS-BSA (PBS + 1% bovine serum albumine) overnight at 4 °C under mild shaking. The next day, worms were centrifuged for 1 minute and then incubated for 24 hours at 4 °C and mild shaking in 500 μL PBS-BSA containing a monoclonal (FG4R), mouse derived anti-flag IgG antibody diluted 1:200 (ab125243, A85282 antibodies.com). The following day, worms were washed again five times in PBS to remove unbound primary antibody. Once again, worms were incubated for 24 hours at 4 °C in 500 μL of PBS-BSA containing DyLight 405 conjugated polyclonal donkey anti-mouse antibodies diluted 1:300 (DyLight™ 405 AffiniPure Donkey Anti-Mouse IgG (H+L), Jackson Immunoresearch). Following another five washes with PBS-BSA, all liquid was completely removed and worms were transferred in ~25 μL to a frosted microscope slide. After adding a drop (~25 μL) of VECTASHIELD® mounting medium, worms were covered with a coverslip and sealed with nail polish. After drying, the antibody staining was examined on an Eclipse Ti-U inverted research microscope (Nikon, Tokyo, Japan) with a 20× and a 40× objective at excitation wavelength 405 nm to visualize antobody specific staining (DyLight405) and 488 nm excitation to visualize pharyngeal GFP expression. Differential interference contrast (DIC) pictures were taken to visualise worm anatomy. Images were taken using VisiView 4.3.0 at 16-bit. ImageJ was used to pseudocolor and merge channels (70). Fluorescent images at 405 nM and 488 nm excitation wavelength were visualized in the blue or green lookup table (LUT) scale provided by ImageJ.

### 4.4 Trashing assay

To prepare stock solutions, IVM and MOX (Sigma-Aldrich) were dissolved in DMSO and frozen at −20 °C. A saturated 40 mM 5-HT stock solution was prepared by dissolving serotonin creatinine sulfate monohydrate (Sigma-Aldrich) in S-medium (67) through vigorous vortexing and was either immediately frozen at −20 °C or used directly.

Bleach synchronized L1 (67) were grown to adult stage on NGM plates (72 h). One day old adults were washed with S-medium by repeatedly allowing worms to sink to the bottom of a 15 ml centrifuge tube, discarding and refilling to remove all bacteria. The arrest of PP caused by the absence of food was confirmed visually after one hour of acclimatisation with an inverted microscope. In each individual experiment, twelve worms (only GFP positive worms in case of transgenic strains) were transferred into 6-well plates (Sarstedt) with 2 ml S-medium final volume containing either no OP50 bacteria (OP50^−^), OP50 bacteria at OD_600_=0.5 (OP50^+^) or 4 mM 5-HT (OP50^−^/5-HT^+^ referred to as 5-HT^+^ in the text for simplicity), the latter two stimulating PP which was visually confirmed before adding drug. Then, a dilution series of both MLs was prepared in DMSO and added to the medium resulting in a final concentration of 1% DMSO. For IVM, the final concentrations for the OP50^−^condition were 0.0, 10.0, 20.0, 30.0, 40.0, 50.0 and 100.0 nM IVM and for the OP50^+^ and 5-HT^+^ conditions 0.0, 1.0, 2.0, 3.0, 4.0, 5.0 and 10 nM IVM. For MOX, final concentrations for all conditions were 0.0, 5.0, 10.0, 12.5, 15.0, 17.5, 20.0, 30, 40, 50 and 100 nM MOX. Plates were sealed with parafilm to avoid concentrations changes through evaporation. Worms were then incubated in the dark at 20 °C and 150 rpm for 18-24 hours. The next day, worms were transferred by pipetting to agar-coated (to prevent worms from sticking to the bottom) petri dishes filled with S-medium without bacteria and were allowed to acclimatise for 1 minute. Then, thrashes of individual worms were counted for one minute on a stereo-fluorescence microscope.

*Caenorhabditis elegans* adult worms which were incubated in S-medium without bacteria did not readily move and even soft touch stimulus and shaking did not induce any movements. However, transfer with a pipette induced strong thrashing, hence this step was essential to detect ML induced paralysis. Likewise, adult worms incubated with a sufficient supply of OP50 bacteria did only thrash occasionally but the transfer of worms by pipetting induced vigorous thrashing.

### 4.5 Statistical Analysis

For each concentration, strain and condition at least 12 worms per day were tested on three separate days (total n = 36). Before log_10_ transformation of drug concentrations, vehicle controls were set to concentrations of 0.1 nM for IVM or MOX. Then, four parameter non-linear logistic regression models and statistical analyses were calculated using GraphPad Prism 8.3.0 (GraphPad Software, San Diego, California USA) constraining the bottom values to ≥0. Visualisation of the half maximal effective concentration (EC_50_) and 95% confidence intervals (CI) values as Forest plots and the corresponding concentration-response curves were made with GraphPad Prism. For the concentration-response curves the negative control concentration was visualised as “0 M no drug” with a break in the x-axis. Statistical differences in EC_50_ values were calculated with the extra-sum-of-squares F test applying the Bonferroni-Holm correction for multiple testing and considering a p-value smaller than 0.05 as significant. For comparison, each *Pun*-PGP-9 expressing transgenic strain was compared to the control strain at the respective condition. The control strain was compared to the WT at the respective condition. EC_50_ values of the WT at the condition without food (OP50^−^) and without food plus 5-HT (5-HT^+^) were compared to the condition with food (OP50^+^). In addition, statistical comparisons of EC_50_ values between MOX and IVM were calculated for the WT. Fold changes were calculated based on EC_50_ values. To examine the impact of condition and transgene expression on motility, the motility responses of all worms from all negative no drug controls were pooled per strain and per condition (n = 72) and a Kruskal-Wallis test with a Dunn’s post-hoc was conducted comparing all conditions in each strain to the WT OP50^−^ condition.

## 7 Data Availability

Plasmids are available upon request and will be deposited on Addgene.org.

## 8 Acknowledgements

We are grateful to Jonathan Ewbank for his guidance with *C. elegans* promotors and for providing the primer sequences for promotor amplification. We thank Jacqueline Hellinga for language proofreading the manuscript.

## 9 Author contributions

APG: Conceptualization, Investigation, Formal analysis, Data Curation and Visualization, Writing – original draft preparation, review and editing, Funding Acquisition

JK: Conceptualization, Data Curation and Visualization, Writing - review and editing, Supervision, Funding Acquisition

CN: Conceptualization, Writing - review and editing, Supervision

CLC: Conceptualization, Writing - review and editing, Supervision, Funding Acquisition

AH: Conceptualization, Writing - review and editing, Supervision

GVS: Conceptualization, Data Curation and Visualization, Writing - review and editing, Supervision, Funding Acquisition

